# Biochemical and Physiological Evidence of Maternal Care in Plants: A Case Study of *Chlorophytum comosum*

**DOI:** 10.64898/2025.12.16.694611

**Authors:** Vineet Soni, Upma Bhatt, Yashwant Sompura

## Abstract

Plants exhibit a wide array of well-characterized survival and reproductive strategies, yet definitive evidence for maternal care has remained absent. Here, we report evidence of maternal care in the plant kingdom using *Chlorophytum comosum* as a model system. Biochemical and physiological analyses were conducted on mother (MRs) and daughter ramets (DRs) of *C. comosum* across four distinct developmental stages of the DRs (S1–S4), ranging from the juvenile stage (S1) to the fully photoautotrophic stage (S4). It was observed that, during the progressive development stages, DRs first develop their photosynthetic potential, followed by the subsequent development of water and mineral uptake capacity. Biochemical and anatomical studies highlighted that stolon acts as a vital life-support conduit, analogous to a placenta, facilitating the translocation of nutrients and water from MRs to DRs. In the present study, the dynamics of sucrose-phosphate synthase, acid invertase, and starch concentrations in MRs and DRs clearly reveal metabolic communication between MRs and DRs and demonstrated the biochemical basis of maternal care in *C. comosum*.

The findings further supported by the stolon severance experiment, in which detached DRs at stage S1 exhibited a 0% survival rate, which increased progressively with developmental stage and reached 100% survival at stage S4. At this stage, when DRs attain full independence, all biological communication between MRs and DRs was terminated. Although DRs achieved full autonomy by Stage S4, MRs maintain stolon-mediated physical connections for approximately two additional weeks, with gradual deterioration at Stage S5 and complete disconnection occurring naturally at Stage S6, indicating post-establishment maternal care. Chlorophyll fluorescence and stolon anatomical analyses during root-prevention and drought-induction experiments revealed that stolon connectivity is maintained when DRs fail to establish roots at Stage S3 or experience drought stress at Stage S4. Under these conditions, MRs continued to provide physiological support via the stolon until successful establishment. Together, these findings demonstrated that *C. comosum* exhibits maternal care until establishment is assured, thereby enhancing survival under environmental stress, and open new avenues for investigating the plant neurobiology and molecular communication underlying maternal care.

## 1. Introduction

Plants represent one of the most mysterious and vital groups of organisms, playing a pivotal role in making Earth a hospitable environment for the survival of other life forms. They provide essential resources such as food, oxygen, shelter, and numerous other necessities, that are essential for maintaining ecological balance. Indeed, if plants were to disappear even for a short period, life on Earth would collapse. Beyond sustaining life, plants actively influence and protect their surrounding environment by shaping microclimates, enhancing soil health, and regulating ecological interactions. These remarkable capacities reflect a form of “care” that plants provide to their environment and to other organisms. However, despite this well-documented ecological “care,” there is currently no evidence that plants extend similar care to their own offspring. This raises a fundamental and largely unexplored question: do plants actively invest in the survival and development of their progeny?

Maternal care, broadly defined as fundamental biological phenomenal investment that enhances the survival and fitness of offspring, represents a central theme in evolutionary biology [1,2]. Maternal care encompasses a diverse range of behaviours and physiological strategies, from nourishment and thermoregulation to protection against predation and pathogens is well known in animals like mammals, birds, and numerous invertebrates [3,4]. These forms of post-embryonic investment are recognized as critical determinants of offspring performance and species persistence. Maternal care is completely unknown in prokaryotes, fungi, and plants, where offspring are typically independent after dispersal and survival relies primarily on intrinsic resilience rather than sustained parental investment. Plants are generally considered devoid of maternal care in this sense. Maternal influence in plants has largely been confined to the provisioning of seeds with reserves, protection through seed coats, and rare instances of vivipary, where germination occurs while seeds remain attached to the maternal plant. While these mechanisms are essential, they represent static and pre-emptive investments rather than dynamic, post-developmental care, which defines maternal behaviour in animals. Thus, the concept of maternal care in plants remains undefined, unrecognized, and untested within physiological or anatomical frameworks.

Stolon-bearing plants offer an exceptional framework to explore potential analogues of maternal support systems. In these systems, developing ramets remain physically and physiologically connected to the mother through stolons that maintain vascular continuity. This anatomical linkage enables the exchange of water, minerals, carbohydrates, and signalling molecules, potentially allowing the maternal plant to influence the establishment and stress resilience of the developing ramet. Yet, despite extensive evidence of stolon-mediated physiological integration, no study to date has interpreted or demonstrated this relationship as a form of maternal care. The few available reports, such as those in *Fragaria* species, merely suggest stolon-assisted nutrient uptake, without examining whether such coordination reflects a dynamic and sustained parental investment [5,6].

In this context, *Chlorophytum comosum* (Asparagaceae), commonly known as spider plant, provides an ideal experimental system to probe this unexplored phenomenon. Its rapid stolon growth, well-defined mother–daughter connectivity, and distinct developmental phases enable precise temporal analysis of resource allocation and metabolic regulation. A systematic investigation of the physiological, biochemical, and anatomical coordination between mother and daughter ramets in *C. comosum* offers a unique opportunity to test whether such integration functions as a plant analogue of maternal care. Establishing this connection would not only challenge the long-standing assumption that plants lack post-embryonic parental strategies but also extend the conceptual boundaries of maternal investment and evolutionary adaptation to the plant kingdom, offering fresh perspectives on plant cooperation, resilience, and developmental plasticity under environmental stress.

This study addresses key questions: How long can mother ramets sustain their daughters? Does maternal support persist under stress or adverse conditions, or do stolons degrade? Which biochemical and physiological parameters define maternal care? Does maternal investment enhance daughter ramet survival, growth synchrony, metabolic stability, and stress tolerance? By interrogating these aspects, we aim to uncover biochemical and physiological basis of maternal care, if exist. This study presents the first integrative evidence of maternal care in plants, combining physiological, biochemical, and anatomical approaches to unravel the dynamic exchange between mother and daughter ramets during development and under stress. Demonstrating this phenomenon could fundamentally reshape our understanding of plant individuality, cooperation, and evolutionary strategies for offspring survival.

## 2 Materials and methods

Spider plants (*Chlorophytum comosum* L.) were chosen as the experimental plant because of their characteristic vegetative propagation through stolons, which produce genetically identical plantlets. Mature and healthy mother plants, grown in pots containing loam soil, were procured from Keshav Nursery, Udaipur, Rajasthan, India in month of June 2025 and transferred at the Plant Bioenergetics and Biotechnology Laboratory, Department of Botany, Mohanlal Sukhadia University, Udaipur, India (24.5854° N, 73.7125° E), where these plants were acclimatized under controlled laboratory conditions (25±2 °C, 12 h light / 12 h dark photoperiod, relative humidity 65 ± 2%) for a period of one month. During this period, the plants exhibited vigorous vegetative growth, and stolon outgrowth. Once the stolons developed daughter ramets at their tips, the experimental design was initiated.

### 2.1 Maternal Care Dynamics Across Sequential Developmental Stages of Daughter Ramets

#### 2.1.1 Experimental Set-up

The older ramets through which stolon raised were referred as mother ramets (MRs) and the newly developed plantlets as daughter ramets (DRs). In order to understand whether plants show maternal care, MRs and DRs of *C. comosum* connected via stolons were studied at following sequential developmental stages (S1–S6) of DRs (Fig. 1):

- **Stage 1 (S1-**Newly formed DRs that were 2 days old): At this stage, the juvenile DRs were still small, fragile, and completely dependent on the MRs for water and nutrients.
- **Stage 2 [S2-**DRs after 8 days of S1 (10 days old)]. By this time, leaves showed further elongation and expansion, but no root initiation was observed. The connection to the MRs remained the sole support of survival of DRs.
- **Stage 3** [**(S3-** DRs after 8 days of S2 (18 days old)]: The most prominent feature of this stage was the onset of root initiation at the basal node of the DRs. Although root initiation was visible but no soil contact was established. At this stage, DRs remain dependent on the MRs for water and nutrients.
- **Stage 4 [(S4-** DRs after 8 days of S3 (26 days old)]: At this stage, the DRs, with well-developed leaves and roots, were established in the soil system and exhibited a higher degree of independence and photoautotrophic potential. This stage represents a transitional phase, where the independent and photoautotrophic DRs remain connected through green and healthy stolons from the MRs.
- **Stage 5 (S5):** Later stage (34 days old) showed the initiation of degradation of stolons.
- **Stage 6 (S6):** The stolons completely degraded (42 days old), which separated MRs and DRs.

**Figure 1.**
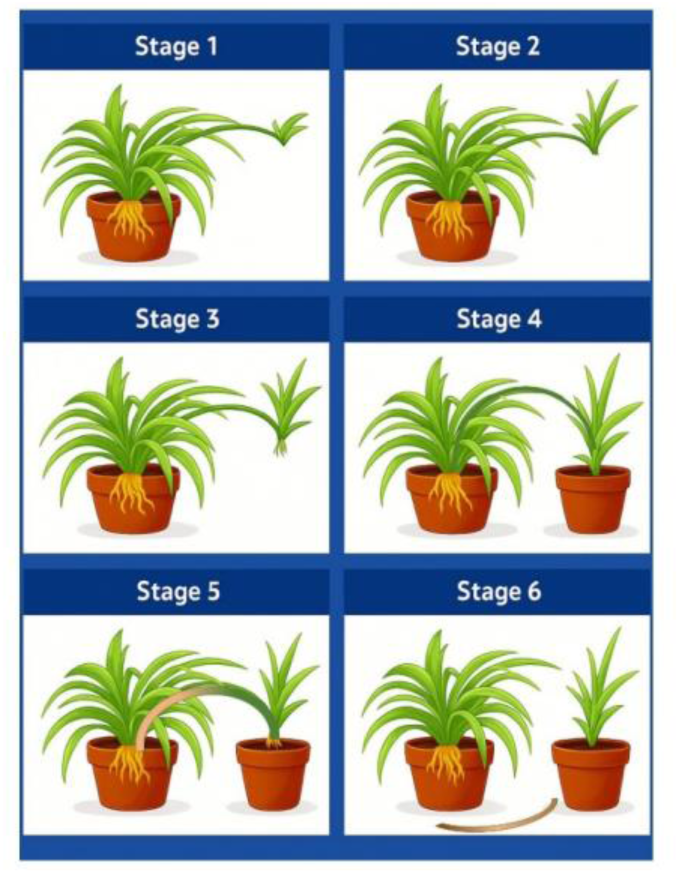
Diagrammatic scheme representing the six defined developmental stages of *Chlorophytum comosum* daughter ramets (DRs) in relation to the mother ramet (MR). Each stage (S1–S6) illustrates characteristic morphological and physiological features, including stolon elongation, root initiation, leaf emergence, and progressive autotrophic independence. The schematic highlights the gradual transition from resource dependence on the mother ramet through the stolon connection (S1–S3) to complete physiological autonomy of the mature DRs and detached stolon (S4–S6).

These developmental stages (S1–S6) provide comparative analyses of anatomical, biochemical and physiological changes during the transition of DRs from juvenile-to-independent growth phases (SF Fig. 1).

#### 2.1.2 Stolon anatomy

Stolons were collected from stages S1 to S6 (Fig. 1). All samples were thoroughly washed with distilled water to remove the surface debris. Free-hand transverse sections were cut from the middle portion of stolon using a sharp razor blade. The thin sections were transferred to a watch glass containing distilled water and then stained with 1% safranin solution for 2–3 minutes. Excess stain was removed by gently washing the sections with distilled water, followed by counterstaining with 0.5% fast green for 30–60 seconds. The stained sections were mounted in 50% glycerol on clean glass slides, covered with a cover slip, and gently placed to avoid air bubbles. The slides were observed under a compound light microscope, and images were captured for detailed examination of anatomical features such as epidermis, cortex and vascular bundles region.

#### 2.1.3 Biochemical analysis

As well-developed rooting and autophototrophic DRs developed at stage S4, all biochemical assays of the DRs, alongside their corresponding MRs, were conducted during their developmental stages from S1 to S4. Leaves were harvested between 11:00 and 12:00 h for bioassays of Ribulose-1,5-bisphosphate Carboxylase (RubisCO), Sucrose-Phosphate Synthase (SPS), Acid Invertase and nitrate reductase (NR) and starch. Prior the bioassay, surface sterilization of leaves was done with 0.1% HgCl_2_ for 3 min followed successive washings with distilled water.

##### Ribulose-1,5-bisphosphate Carboxylase

RubisCO activity was determined in leaf tissues of MRs and DRs by following the modified method of [7]. Leaves were homogenized in ice-cold extraction buffer (100 mM Bicine, pH 7.8; 5 mM EDTA; 0.75% PEG; 14 mM β-mercaptoethanol; 1% Tween-80; 1.5% PVP), and centrifuged at 13,000 × g for 40 s. The supernatant was assayed at 25 °C by coupling RuBP carboxylation to NADH oxidation, monitored at 340 nm (Shimadzu UV-160). Initial activity was measured immediately after adding RuBP, while total activity was measured after full activation of RuBisCO by a 15 min pre-incubation with assay solution. The activation state was expressed as the ratio of initial to total activity, allowing comparison between MRs and DRs.

##### Sucrose-Phosphate Synthase

SPS activity was measured in the leaf tissues of MRs and DRs. The assay followed the protocol of [8] 1 gm leaves were harvested from each sample. Leaf tissue was homogenized in pre-chilled extraction buffer. The homogenate was centrifuged at 15,000 × g for 15 min at 4°C, and the supernatant was collected as the crude enzyme extract.

SPS activity was assayed by incubating aliquots of the enzyme extract with a reaction mixture containing UDP-glucose, fructose-6-phosphate, and Mg²⁺ ions for 20 min under linear reaction conditions. The reaction was terminated by adding the stop reagent, and sucrose formation was quantified colorimetrically at 540 nm using the anthrone method. Enzyme activity was expressed as μmol sucrose formed per minute per gram fresh weight (μmol min⁻¹ g⁻¹ FW).

##### Acid Invertase

Acid invertase activity was determined by quantifying glucose released from sucrose hydrolysis, following [9]. A glucose standard curve was prepared in the range of 0.2–2.0 mg mL⁻¹ in distilled water. The enzyme assay was performed in a 1.0 mL reaction mixture containing 0.4 mL acetate buffer (0.2 M, pH 4.8), 0.2 mL sucrose solution (0.4 M), and 0.4 mL of enzyme extract. The mixture was incubated at 30 °C for 60 min. The reaction was terminated by adding 1.0 mL of dinitrosalicylic acid (DNS) reagent, followed by heating in a boiling water bath for 5 min. Samples were rapidly cooled under running tap water, and 8.0 mL of distilled water was added to adjust the final volume to 10 mL. Absorbance was recorded at 560 nm against a blank in which DNS reagent was added at zero time (prior to incubation). Enzyme activity was expressed as the amount of glucose released (mg) per hour per mg of protein, based on the glucose standard curve.

##### Nitrate reductase

*In vivo* nitrate reductase activity was assayed in leaf tissues of MRs and DRs following [10] Leaf segments (600 mg fresh weight) were incubated in 10 mL assay solution containing 100 mM phosphate buffer (pH 7.5), 30 mM KNO₃, and 5% propanol under dim light at 35 °C for 30 min with gentle shaking (80 rpm). Samples were boiled at 100 °C for 5 min prior to incubation to inactivate enzyme activity. After incubation, reactions were terminated by boiling, cooled, and nitrite (NO₂^-^) produced was quantified colorimetrically at 540 nm using sulfanilamide (1% in HCl) and N-(1-naphthyl)-ethylenediamine dihydrochloride (0.02%) as Griess reagents. A KNO₂ standard curve (25 μM) was used for calibration, and enzyme activity was expressed as μmol NO₂⁻produced g⁻¹ FW h⁻¹.

##### Starch

Starch content was estimated following the acid hydrolysis method of [11]. Residual soluble sugars were removed from the ethanol extract residue by washing with 80% ethanol. The residue was then extracted with 52% perchloric acid in repeated steps to a final volume of 100 mL. A diluted aliquot (0.5 mL) was mixed with distilled water and anthrone reagent (2g anthrone in 1 L of 96% H_2_SO_4_) in an ice bath, heated at 100 °C for 8 min, and cooled rapidly. Absorbance was measured at 630 nm, and starch content was quantified using a glucose standard curve.

#### 2.1.4 Biophysical analysis

##### Chlorophyll *a* fluorescence kinetics

Chlorophyll *a* fluorescence O-J-I-P transients were recorded using a Plant Efficiency Analyzer (Hansatech Instruments, Kings Lynn, Norfolk, U.K.). Whole plant samples were dark-adapted for 1 h, and the measured leaf areas were further darkened in clips for 1 min before measurement. A high-intensity LED array (peak wavelength 650 nm, 3000 µmol m⁻² s⁻¹) illuminated a 4 mm leaf area, ensuring complete closure of Photosystem II reaction centers. Transients were recorded for 1 s, and fluorescence intensities were obtained at 50 µs, 300 µs, 2 ms, 30 ms and at the P peak [12].

##### Specific, phenomenological and quantum yields and performance index

Photosynthetic parameters such as maximum fluorescence (Fm), specific energy fluxes (ABS/RC, TR/RC, ET/RC, DI/RC), phenomenological fluxes (ABS/CS, TR/CS, ET/CS, DI/CS), density of functional PSII reaction centers (RC/CS), quantum efficiencies of photosynthesis (ϕP_o_) and electron transport (ϕE_o_), and performance index (PIcs) were subsequently calculated as per the equations of JIP test [13] (SF Table 1).

**Table 1.** Changes in JIP test parameters in *Chlorophytum comosum* under root prevention and drought induction experiments. Values represent mean ± standard error (SE) of three biological replicates. MR and DR denote mother and daughter ramets, respectively, while S3a, S4a, and S5 indicate specific developmental stages or days after treatment. Different superscript letters (a–c) within a row indicate statistically significant differences among treatments at *p* < 0.05 (estimated visually; actual significance to be confirmed by one-way ANOVA followed by Tukey’s post hoc test).

| Parameters | MR (S5) | MR (S3a) | DR (S3a) | MR (S4a) | DR (S4a) |
| --- | --- | --- | --- | --- | --- |
| ***F<sub>m</sub>*** | 1066.67 ± 53.41<sup>a</sup> | 1094.67 ± 13.43<sup>a</sup> | 902.33 ± 12.42<sup>b</sup> | 1072.67 ± 38.55<sup>a</sup> | 886.33 ± 88.07<sup>b</sup> |
| ***PHI(Po)*** | 0.75 ± 0.03<sup>a</sup> | 0.80 ± 0.00<sup>a</sup> | 0.65 ± 0.16<sup>ab</sup> | 0.79 ± 0.01<sup>a</sup> | 0.39 ± 0.07<sup>b</sup> |
| ***PHI(Eo)*** | 0.48 ± 0.00<sup>b</sup> | 0.56 ± 0.01<sup>a</sup> | 0.35 ± 0.17<sup>c</sup> | 0.58 ± 0.02<sup>a</sup> | 0.18 ± 0.06<sup>c</sup> |
| ***RC/CS<sub>m</sub>*** | 623.72 ± 85.73<sup>b</sup> | 944.45 ± 54.07<sup>a</sup> | 468.22 ± 228.57<sup>b</sup> | 829.09 ± 73.98<sup>a</sup> | 172.29 ± 70.48<sup>c</sup> |
| ***ABS/CS<sub>m</sub>*** | 1066.67 ± 53.41<sup>a</sup> | 1094.67 ± 13.43<sup>a</sup> | 902.33 ± 12.42<sup>b</sup> | 1072.67 ± 38.55<sup>a</sup> | 886.33 ± 88.07<sup>b</sup> |
| ***TRo/CS<sub>m</sub>*** | 804.66 ± 66.74<sup>ab</sup> | 879.64 ± 15.96<sup>a</sup> | 589.67 ± 148.01<sup>b</sup> | 852.32 ± 26.11<sup>a</sup> | 350.65 ± 95.63<sup>c</sup> |
| ***ETo/CS<sub>m</sub>*** | 515.68 ± 30.92<sup>b</sup> | 617.65 ± 2.06<sup>a</sup> | 316.02 ± 153.11<sup>c</sup> | 623.67 ± 41.29<sup>a</sup> | 158.99 ± 65.38<sup>c</sup> |
| ***DIo/CS<sub>m</sub>*** | 262.00 ± 14.97<sup>b</sup> | 215.02 ± 2.66<sup>b</sup> | 312.66 ± 135.64<sup>b</sup> | 220.34 ± 14.55<sup>b</sup> | 535.68 ± 36.78<sup>a</sup> |
| ***PIcs*** | 34539.03 ± 6873.32<sup>b</sup> | 91140.61 ± 1987.55<sup>a</sup> | 17690.80 ± 15225.68<sup>b</sup> | 88638.67 ± 16911.72<sup>a</sup> | 11060.21 ± 1013.16<sup>c</sup> |

### 2.2 Maternal Dependency of Daughter Ramets Under Stolon Severance, Root Restriction, and Drought Stress

#### 2.2.1 Stolon Severance experiment

DRs at different developmental stages (S1–S4) were carefully excised from the stolons and transplanted into pots filled with loam soil to analyse RWC and survival rate.

##### Relative Water Content (RWC) Measurement

Relative water content (RWC) of DRs was determined following the method of [12] RWC was monitored at 2, 4, 6, and 8 days post-transplantation to evaluate their water status relative to the attached MRs. For each sampling point, leaves were excised, and fresh weight (FW) was immediately recorded using a precision balance. Samples were then floated on distilled water under controlled light conditions for 4 h to attain full hydration, after which the turgid weight (TW) was measured. Subsequently, the samples were oven-dried at 80 °C for 24 h to obtain dry weight (DW). RWC (%) was calculated using the following formula:

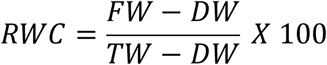

##### Survival Rate Measurement

Survival rate of DRs (S1–S4) was assessed after 8 days of detachment from the MRs. For this, all DRs were excised and transplanted into independent soil-filled pots under controlled growth conditions, similar to those used for RWC determination. The plants were monitored daily for visual symptoms of survival or mortality (leaf wilting, chlorosis, necrosis, or complete drying). At the end of the 8th day, the number of surviving DRs (those maintaining green leaves, turgidity, and active growth) was recorded in comparison to the total number of transplanted ramets. Survival rate (%) was calculated using the following formula:

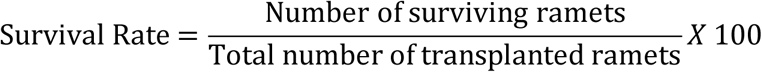

#### 2.2.2 Root prevention experiment

In the experiment, rooting of DRs, maintained at Stage 3, was prevented from contacting the soil by tightly placing a fine nylon mesh (pore size ∼100–200 µm) over the pots. This barrier effectively prevented direct root–soil contact, thereby blocking the transition from Stage 3 to Stage 4 and inhibiting the autonomous establishment of the DRs (Fig. 2a). Importantly, the stolon connection to the MRs remained intact, ensuring vascular continuity while eliminating the possibility of independent soil-derived nutrition. Comparative analyses were performed with reference to S5 (34-day-old ramets), as the treated plants advanced developmentally to the S5 stage following the treatment period.

**Figure 2.**
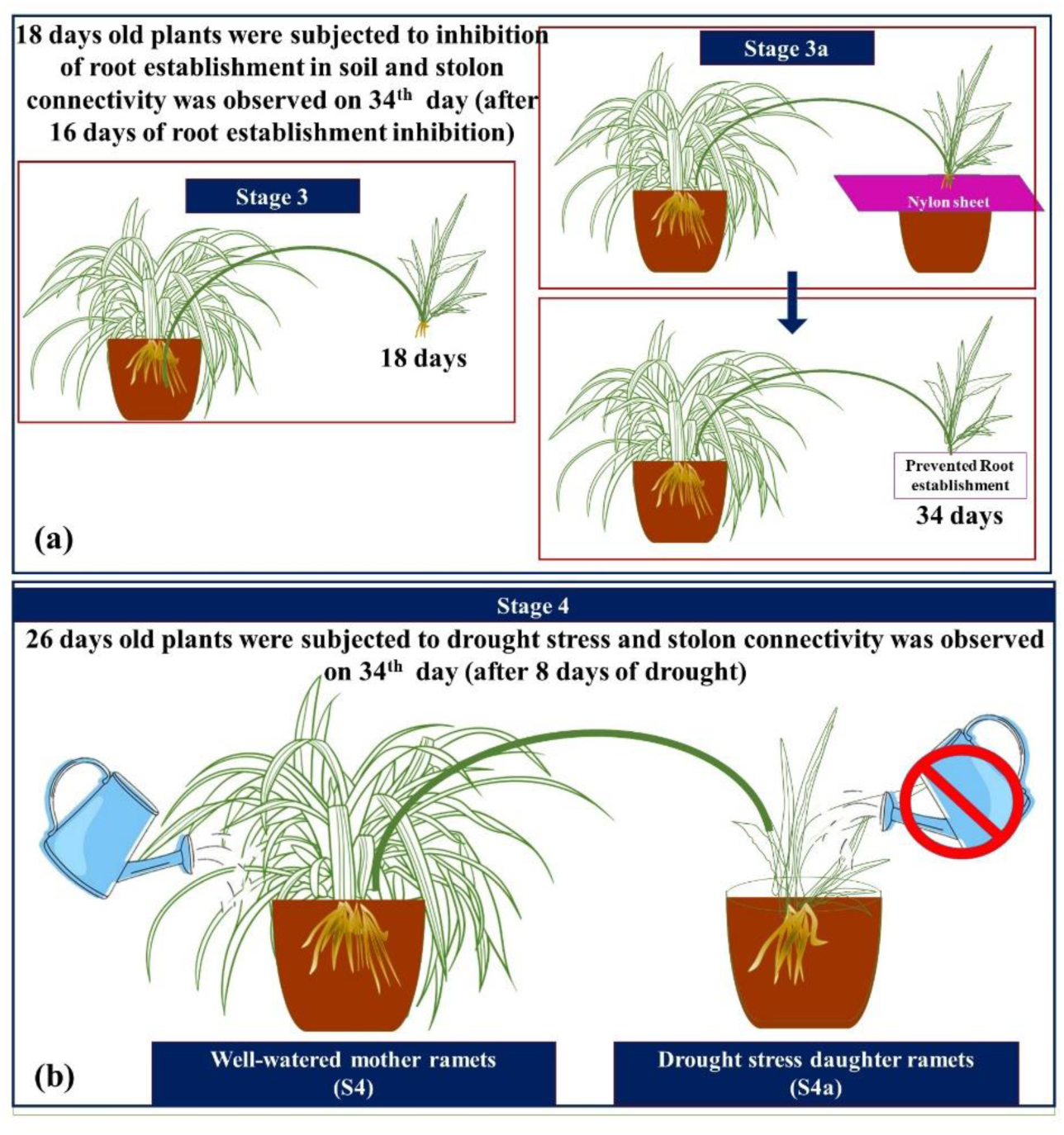
Schematic representation of the experimental design used for studying physiological integration and stress response in *Chlorophytum comosum*. *(a) Root prevention experiment* — illustrating the setup in which stolon-mediated connection was maintained between mother ramets (MRs) and daughter ramets (DRs), while root formation of DRs was physically restricted to assess the contribution of maternal support. (b) Drought-induced experiment — depicting the controlled water deficit treatment applied to DRs to evaluate the role of maternal linkage in modulating drought tolerance mechanisms. Both setups were maintained under identical environmental conditions, and corresponding measurements were taken at defined developmental phases.

##### Stolon anatomy

Stolon anatomy at S3 was evaluated by preventing root establishment of DRs of S3. All plants were maintained under well-watered, controlled conditions (25 ± 2 °C, 50–60% RH, 12 h photoperiod, light ≈150–200 μmol m⁻² s⁻¹). At the beginning of the root prevention experiment, the DRs were 18 days old. The experiment continued for 16 days, making the plants 34 days old by the end of the treatment period. Therefore, a comparative analysis was performed between 18-day-old and 34-day-old DRs. Transverse sections were prepared with a fresh razor, stain with 1% safranin for 2–3 min, rinse in distilled water, counterstain with 0.5% fast green for 30–60 s, mount in 50% glycerol, and observe under a compound microscope. Photomicrographs for each sample were captured.

##### Chlorophyll *a* fluorescence kinetics and JIP test

Chlorophyll *a* fluorescence and all previously described JIP-test parameters were analyzed in MRs and DRs after 16 days of the root-prevention experiment (i.e., 34 days for both DRs and MRs), and the results were compared with plants at the S5 developmental stage, which are naturally 34 days old.

#### 2.2.3 Drought induction experiment (S4 stage)

The experiments were conducted at the S4 stage, when DRs had already established roots in soil but were still connected to the MRs via stolons. At the onset of S4 (26 days old), drought stress was imposed by ceasing water supply to the pots containing DRs, while the corresponding MRs were maintained under well-watered conditions (Fig. 2b). Control pairs received regular irrigation for both mother and daughter pots. Plants were grown under controlled environment (25 ± 2 °C, 50–60% RH, 12 h photoperiod, light ≈150–200 μmol m⁻² s⁻¹). Observations were made after 8 days of drought treatment (34 days old DRs). Comparative analyses were performed with reference to S5 (34-day-old ramets), as the treated plants advanced developmentally to the S5 stage following the treatment period.

##### Stolon anatomy

To evaluate stolon-mediated support under stress were measured. At day 34, transverse sections of stolons were also prepared with a fresh razor, stained with 1% safranin (2–3 min), rinsed, counterstained with 0.5% fast green (30–60 s), mounted in 50% glycerol, and observed under a compound microscope. Vascular bundle condition (intact vs. collapsed) and tissue degradation were documented, and representative photomicrographs were captured.

##### Chlorophyll *a* fluorescence kinetics and JIP test

Chlorophyll *a* fluorescence and all JIP-test parameters were measured in MRs and DRs after 8 days of drought induction (34 days old), and compared with naturally 34-day-old plants at the S5 developmental stage.

### 2.3 Statistical analysis

All experimental data were expressed as mean ± standard deviation (SD) of nine independent biological replicates for each developmental group of *C. comosum* (Spider plant). Statistical analyses were performed to evaluate the significance of variations among the treatment groups. One-way analysis of variance (ANOVA) was carried out, followed by post-hoc comparison tests to determine significant differences at *p* < 0.05.

## 3. Results

### 3.1 Maternal Care Dynamics Across Sequential Developmental Stages of Daughter Ramets

#### a) Stolon anatomy across DRs developmental stages

Microscopic examination of cross sections across successive stages of the mother–daughter junction revealed clear histological transitions. At S1, the stolon consisted of a single-layered epidermis with a thin cuticle, a narrow chlorenchymatous zone, and a parenchymatous cortex, while the vascular region remained poorly differentiated with only a few procambial traces (Fig. 3a). In S2, the epidermis was intact, the cortex enlarged with compact parenchyma, and vascular strands exhibited clearer xylem and phloem differentiation (Fig. 3b). By S3, the cortex thickened and vascular bundles became well organized, with xylem elements showing lignification and phloem cells more distinctly defined (Fig. 3c). At S4, a continuous epidermis, multi-layered parenchymatous cortex, and highly differentiated vascular bundles characterized a fully matured stolon (Fig. 3d). In S5, the onset of senescence was evident, with a wrinkled epidermis, cortical shrinkage, partial disintegration, and collapsing vascular elements, although some xylem vessels retained lignified walls (Fig. 3e). Finally, S6 exhibited advanced tissue degeneration, marked by epidermal collapse, complete cortical disintegration, and disorganized vascular bundles, leaving behind thick-walled, lignified sclerenchymatous remnants of the original vascular structure (Fig. 3f).

**Figure 3.**
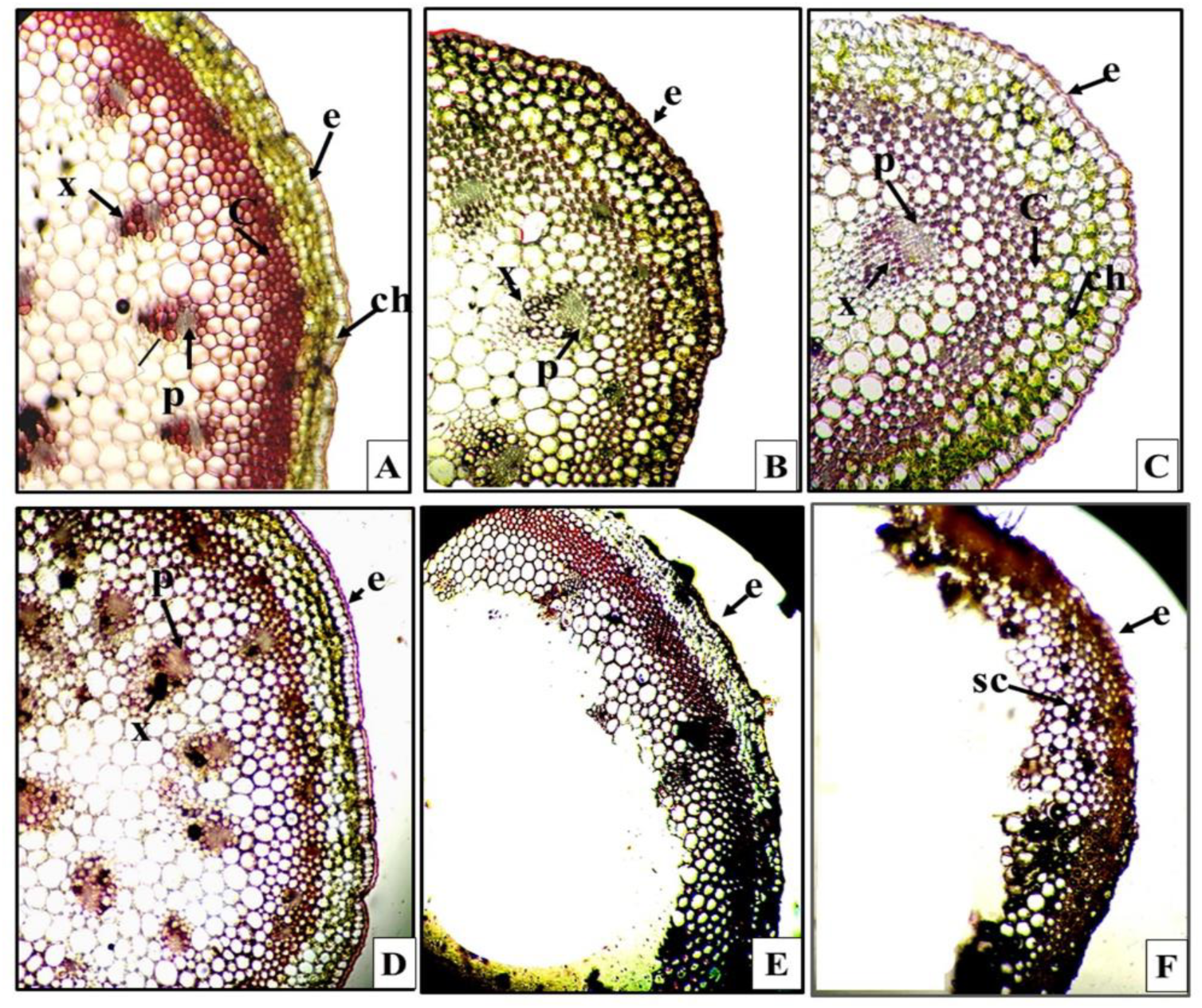
Photomicrographic illustration showing transverse sections of stolons at successive developmental stages (S1 to S6) of the daughter ramet in *Chlorophytum comosum*. *(A)* Immature daughter ramet exhibiting primary tissue differentiation with distinct epidermis (e), collenchyma (c), chlorenchyma (ch), phloem (p), and xylem (x). *(B)* Root initiation stage showing organized vascular bundles with clearly defined xylem (x) and phloem (p). *(C)* Soil-established daughter ramet still connected to the mother ramet through the stolon, displaying well-developed vascular tissues (x, p). *(D)* Onset of stolon degradation characterized by initial tissue disintegration. *(E)* Advanced degradation stage showing complete disruption of tissue organization. *(F)* Terminal stage with a dried stolon exhibiting lignified sclerenchymatous cells (sc).

#### b) Biochemical analysis

##### RuBisCO activity

RuBisCO activity displayed a distinct stage-dependent pattern between MRs and DRs. In S1 phase, DRs exhibited minimal RuBisCO activity (15 µmol g⁻¹ FW h⁻¹), indicating their strong dependence on maternal support, while MRs recorded the maximum activity (78.5 µmol g⁻¹ FW h⁻¹). As development advanced, a gradual increase in RuBisCO activity was observed in DRs. At S3, DRs displayed a further increment, while MRs showed a slight reduction, thereby narrowing the difference between the two (Fig. 4a). At S4, RuBisCO activity in DRs reached levels comparable to the MRs, suggesting a shift toward photosynthetic independence.

**Figure 4.**
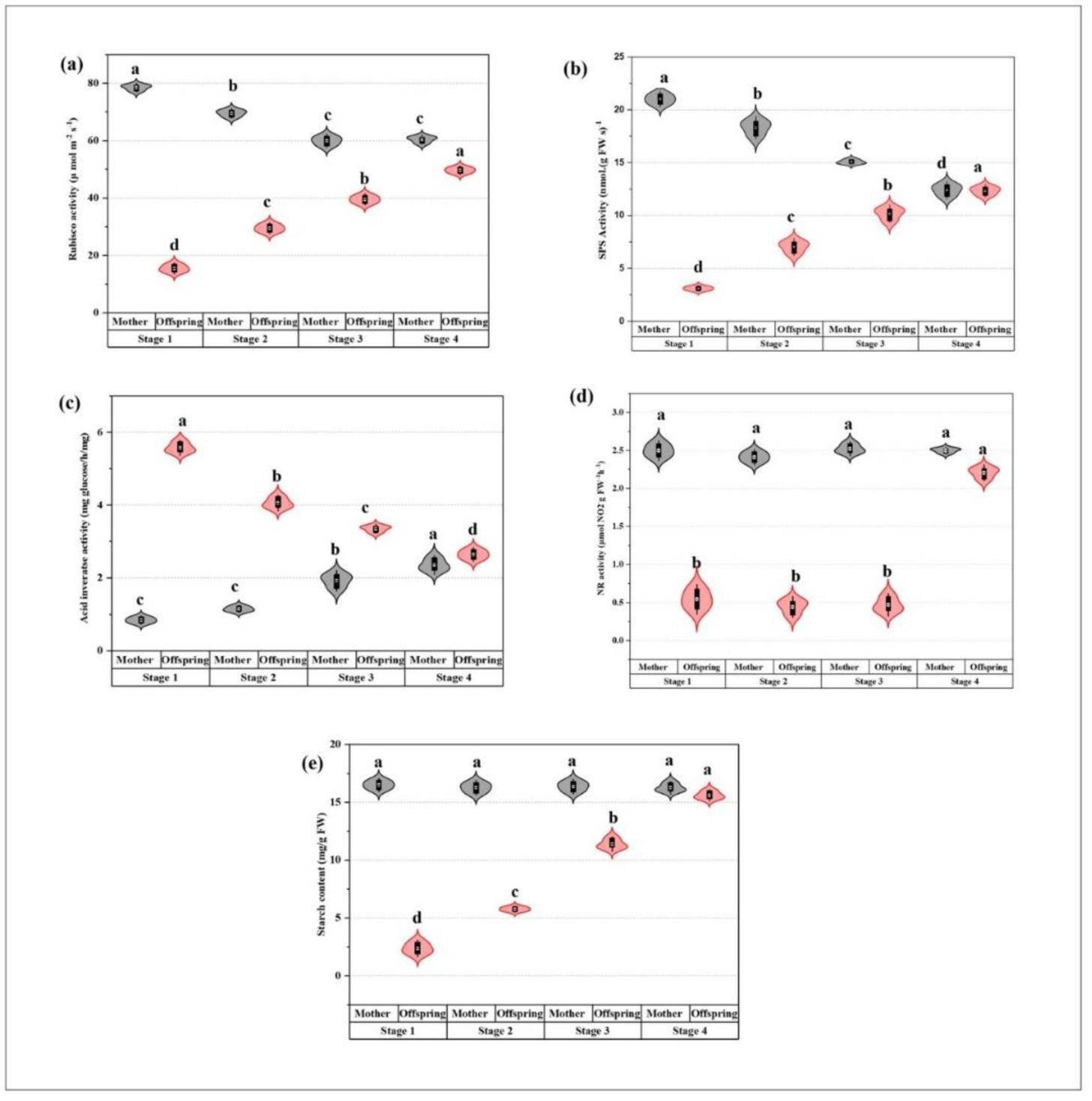
Changes in key metabolic parameters during sequential developmental stages (S1–S4) in mother ramets (MRs) and daughter ramets (DRs) of *Chlorophytum comosum*. Activities of (a) Ribulose-1,5-bisphosphate carboxylase, (b) Sucrose phosphate synthase, (c) Acid invertase, (d) Nitrate reductase, and (e) starch content. Different letters indicate significant differences at *p* ≤ 0.05.

##### SPS (Sucrose Phosphate Synthase) activity

SPS activity reflecting the gradual shift from maternal support to autonomous metabolic functioning in DRs. At S1, offspring exhibited only 14.8% of maternal SPS activity. By S2, offspring SPS activity rose to 38% of the mother’s, indicating the initiation of their own metabolic contribution, although maternal dominance persisted. A marked shift occurred at S3, where offspring reached 67.5% of maternal activity, suggesting a substantial enhancement of their sucrose biosynthetic capacity. By S4, DRs attained 99% of the MRs SPS activity, effectively converging with maternal levels (Fig. 4b). The concurrent decline in maternal SPS activity reflects a strategic transition of energetic responsibility from MRs to DRs during later developmental phases.

##### Acid invertase activity

Acid invertase activity exhibited contrasting trends between MRs and DRs, highlighting differential roles during development. In MRs, invertase activity was initially very low at S1 (0.84 µmol g⁻¹ FW h⁻¹) but showed a steady, stage-wise increase, reaching 2.36 µmol g⁻¹ FW h⁻¹ at S4. In contrast, DRs displayed markedly higher acid invertase activity at the onset, recording 5.57 µmol g⁻¹ FW h⁻¹ at S1 up to sixfold greater than their mothers (Fig. 4c). However, unlike the MRs, DRs showed a progressive decline in activity over subsequent stages, and stabilizing near maternal levels (2.64 µmol g⁻¹ FW h⁻¹) by S4.

##### NR (Nitrate Reductase) activity

NR activity remained consistently high in MRs across all developmental stages (2.41–2.52 µmol NO₂⁻ g⁻¹ FW h⁻¹), showing no significant variation between stages 1 to 4. In contrast, the DRs exhibited very low NR activity during the early developmental stages (0.44–0.54 µmol NO₂⁻ g⁻¹ FW h⁻¹ at S1–3). A marked increase was observed at S4, where DRs established in soil and their NR activity (2.20 µmol NO₂⁻ g⁻¹ FW h⁻¹) approached that of the MRs (Fig. 4d). This pattern indicates that NR activity in DRs is negligible prior to root establishment and becomes strongly induced once independent nitrate uptake from soil is possible.

##### Starch content

Starch content varied significantly across developmental stages. Starch content in MRs remained almost unchanged across all stages (16.26–16.50 mg g⁻¹ FW), showing less than 2% variation overall. In contrast, DRs displayed a sharp developmental increase in starch levels. At S1, DRs had extremely low starch levels, where as S2 and S3 showed a gradual increase in starch content was observed up to 144 and 381%, bringing them closer to maternal values (Fig. 4e). By S4, DRs accumulated 15.60 mg g⁻¹ FW starch and reached up to 96% of the starch level measured in MRs.

#### c) Biophysical analysis

##### Chlorophyll *a* fluorescence kinetics

The fast chlorophyll *a* fluorescence induction curves (OJIP transients) demonstrated distinct differences between MRs and DRs. MRs displayed a typical O–J–I–P rise with high fluorescence values. S1 DRs showed markedly suppressed transients, with much lower O, J, I, and P levels compared to the MRs, reflecting poor PSII performance (Fig. 5a). S2 offspring indicated partial recovery, while S3 ramets revealed strong recovery, with I and P steps nearly overlapping maternal values. The fluorescence curve became almost identical to mothers at S4 phase of DRs. Minimal Fluorescence (Fo) was lowest in S1 DRs and showed intermediate recovery during S2 and S3 phase (Fig. 5b). Maximum fluorescence (F_m_) shown a similar pattern, with MRs and advanced DRs stages (S3–S4) displaying comparable values, whereas S1 DRs showed markedly reduced levels (Fig. 5c).

**Figure 5.**
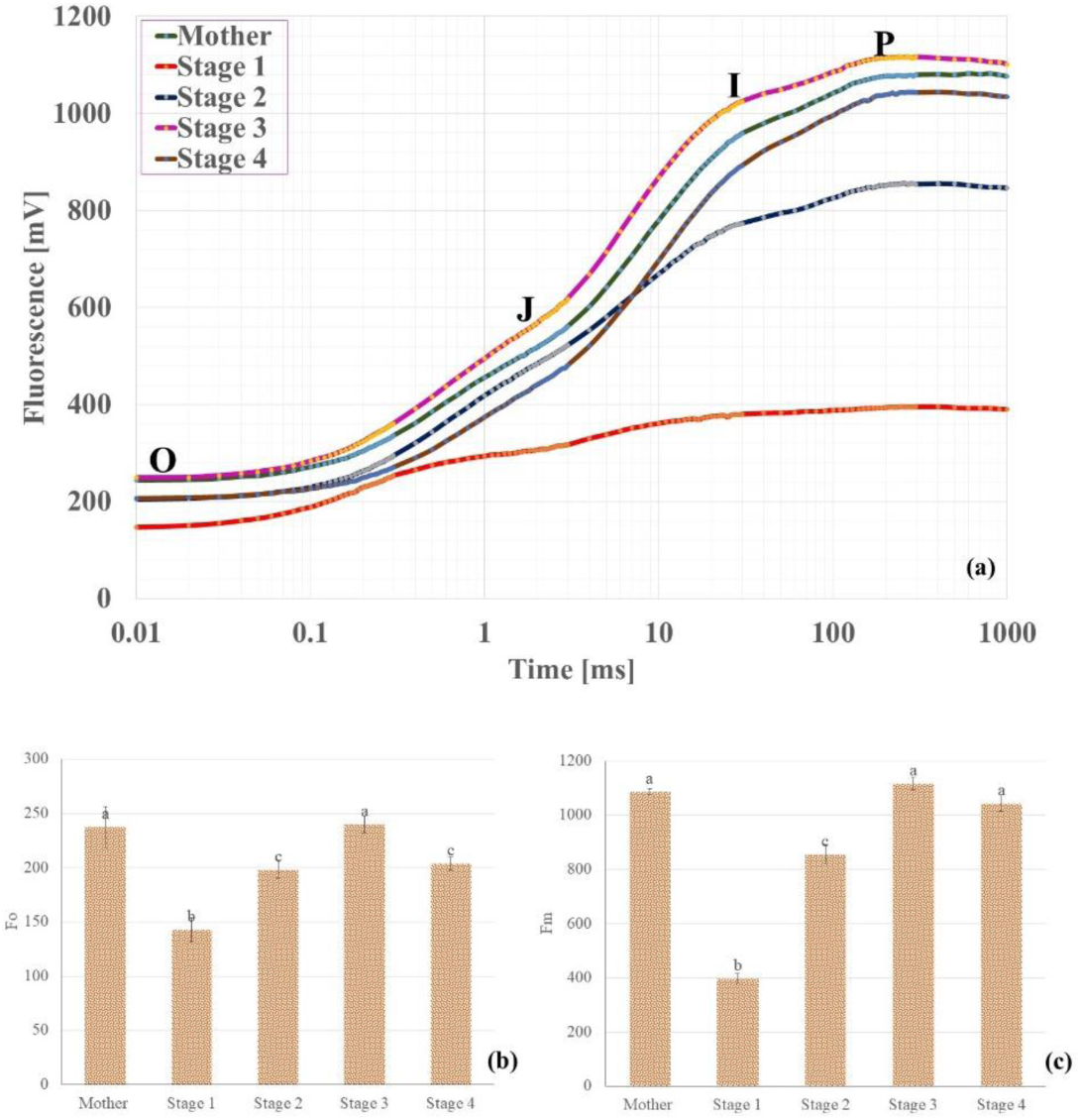
Alteration in OJIP transient curve during sequential developmental stages (S1–S4) in mother ramets (MR) and daughter ramets (DRs) of *Chlorophytum comosum* (a). Bar graphs (b) and (c) depict minimal fluorescence (F_0_) and maximal fluorescence (F_m_), respectively. Different letters indicate significant differences at *p* ≤ 0.05.

##### Specific energy fluxes per reaction center

The absorption flux per reaction center (ABS/RC) was significantly higher in stage S1 DRs compared to MRs (Fig. 6a). However, ABS/RC declined in subsequent stages and reached levels comparable to those of MRs. Similarly, the trapped energy flux (TR_0_/RC) was also elevated in S1 DRs, suggesting greater excitation pressure per PSII unit (Fig. 6b). This parameter normalized by S2 and remained comparable to MRs throughout the further development of DRs. In contrast, the electron transport flux (ET_0_/RC) did not vary significantly across stages, indicating that downstream electron transport capacity remained relatively stable once excitation energy was trapped (Fig. 6c). Dissipated energy per reaction center (DI_0_/RC) was markedly higher in S1 DRs, but values converged with those of the MRs at later stages (Fig. 6d).

**Figure 6.**
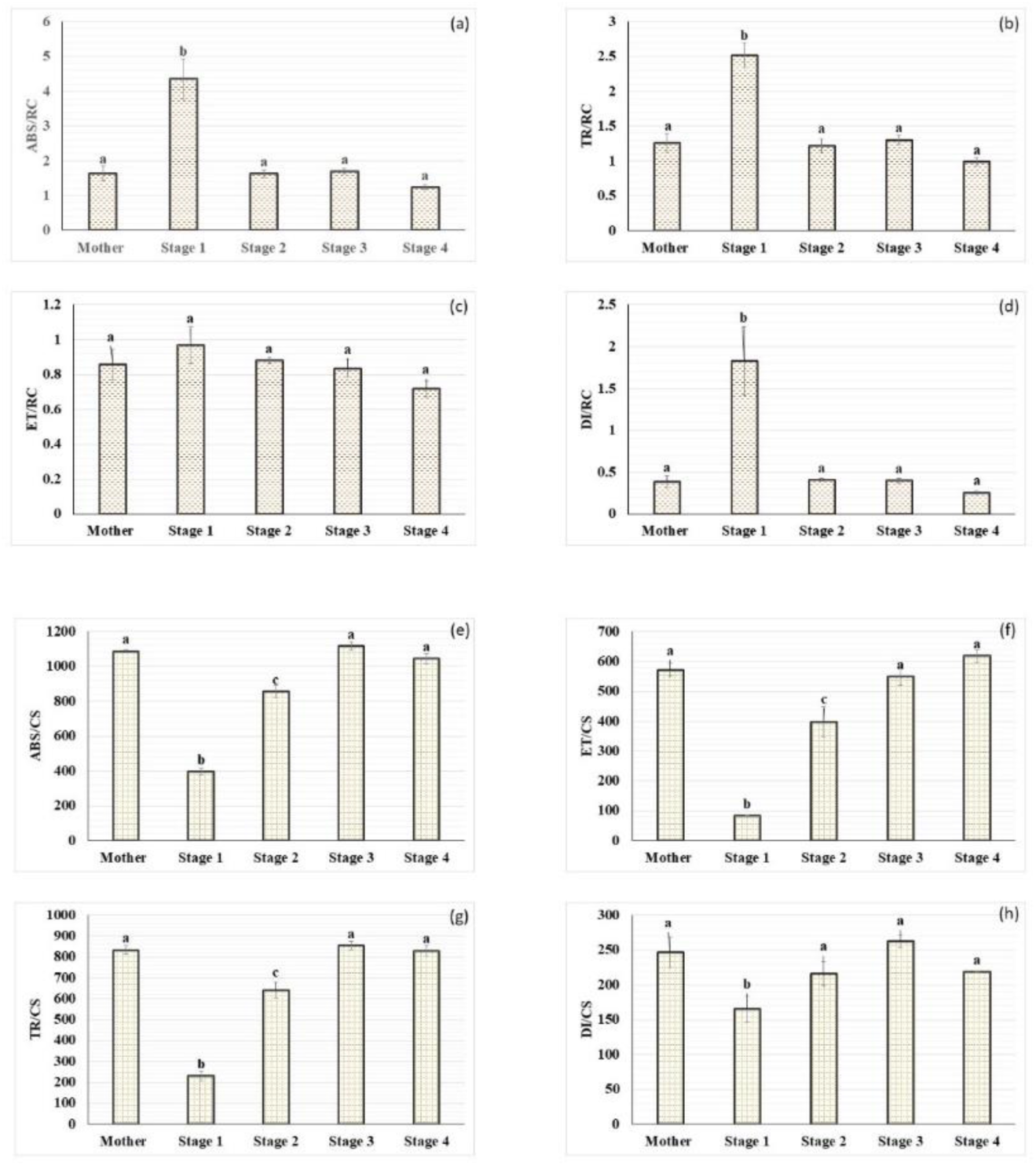
Specific and phenomenological energy fluxes of PSII in mother ramets (MR) and daughter ramets (DRs) of *Chlorophytum comosum* during sequential developmental stages (S1–S4). (a) absorption flux (ABS/RC), (b) trapped energy flux (TR_0_/RC), (c) electron transport flux (ET_0_/RC), and (d) dissipated energy flux (DI_0_/RC). While (e–h) show phenomenological fluxes per cross-section: (e) absorption flux (ABS/CS), (f) trapped energy flux (TR_0_/CS), (g) electron transport flux (ET_0_/CS), and (h) dissipated energy flux (DI_0_/CS). Different letters indicate significant differences at *p* ≤ 0.05.

##### Phenomenological energy fluxes per cross-section

The absorption flux per cross section (ABS/CS) was higher in MRs (Fig. 6e). In contrast, DRs showed the lowest ABS/CS values at S1, which gradually increased and reached maternal levels by S3, continuing similarly through S4. A similar trend was observed for trapped energy flux per cross section (TR_0_/CS), where S1 DRs showed minimal values (∼200), while subsequent stages recovered progressively to maternal levels (Fig. 6f). Electron transport per cross section (ET_0_/CS) was also extremely low in S1 DRs compared to MRs, but values increased steadily and reached maternal levels by S3–S4 (Fig. 6g). Interestingly, dissipated energy per cross section (DI_0_/CS) remained lowest in S1 DRs relative to MRs and later developmental stages, indicating that the limited overall absorption capacity restricted energy dissipation per unit cross section during the early phase (SF Fig. 2).

##### Density of functional reaction center, PSII performance and efficiency parameters

The density of active reaction centers per cross-section (RC/CS) revealed sharp differences between MRs and DRs across developmental stages (Fig. 7a). In contrast, S1 DRs exhibited drastically reduced RC/CS, reflecting a significant loss of functional PSII units during early establishment. However, S2 DRs showed a strong recovery, and by S3 values nearly matched the mother. S4 DRs surpassed maternal values, reaching 841.6, suggesting that mature DRs develop a higher density of active PSII centers compared to mothers, signifying a transition toward photosynthetic independence.

**Figure 7.**
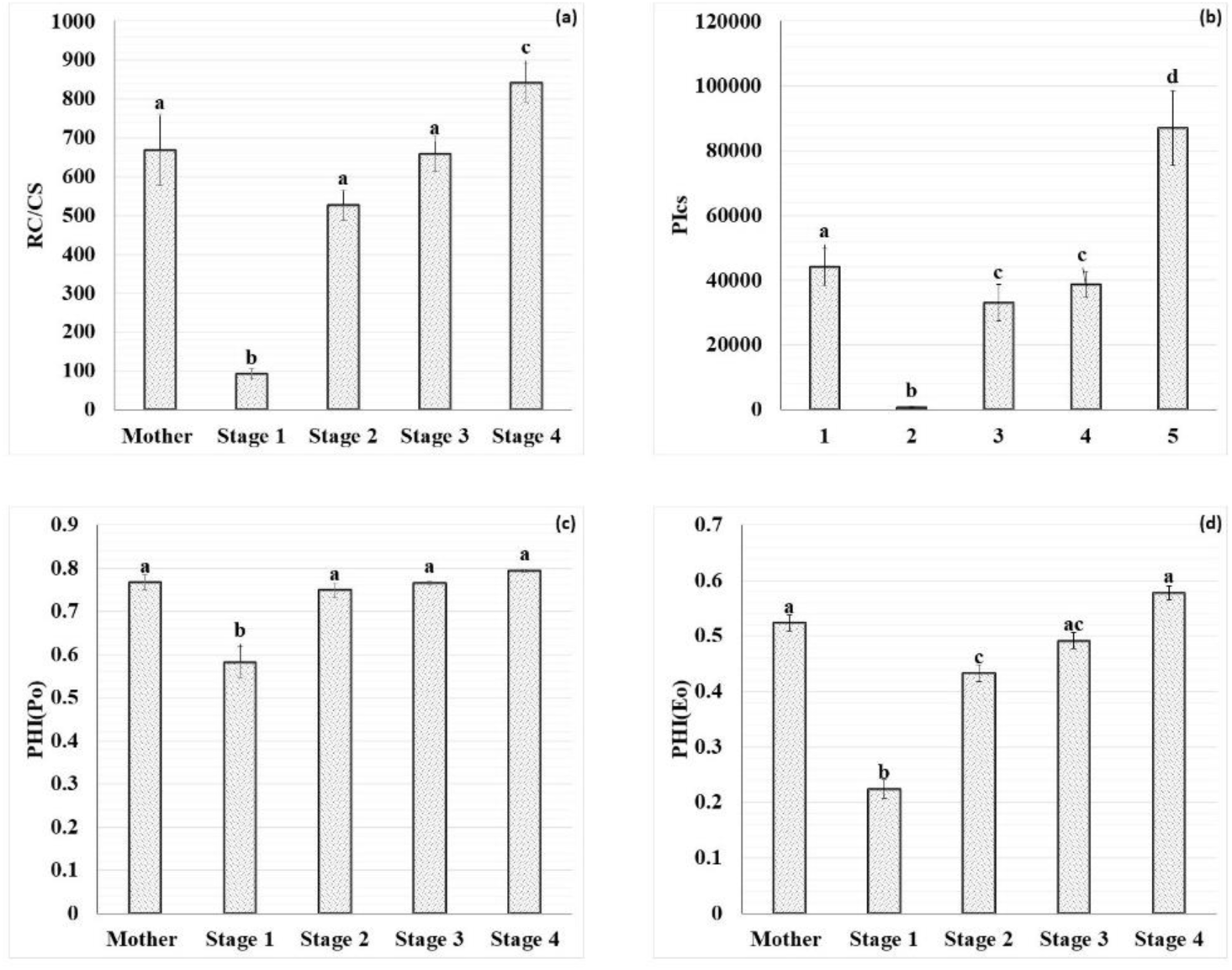
Variations in (a) Density of active reaction centers per cross section (RC/CS), (b) Performance index per cross section (PIcs), (c) Maximum quantum yield of primary photochemistry (ΦP_0_), and (d) Quantum yield for electron transport beyond QA⁻ (ΦE_0_) at different development stages (S1–S4). Different letters indicate significant differences at *p* ≤ 0.05.

The performance index based on energy conservation from photon absorption to electron transport (PIcs) further highlighted the developmental differences. MRs exhibited high PIcs values, while S1 DRs showed an extreme collapse up to 820, confirming severe PSII inefficiency at early development. DRs of S2 and S3 phases demonstrated partial recovery, though still significantly lower than mothers. By S4, PIcs values showed a dramatic increase, exceeding maternal levels and reflecting enhanced structural and functional reorganization of PSII in established DRs (Fig. 7b).

Quantum yield parameters exhibited a more gradual recovery trend. The maximum quantum yield of primary photochemistry (ΦPo) was stable across mothers and offspring except for S1 (Fig. 7c). S1 offspring showed a significant reduction up to 0.58, indicating impaired efficiency of PSII charge separation. Similarly, the quantum yield of electron transport (ΦEo) followed a comparable pattern. MRs maintained high values 0.52, while S1 DRs showed the lowest values 0.22, confirming a major blockage in electron transport beyond Q_A_ (Fig. 7d). DRs of S2 exhibited moderate recovery, while S3 and S4 DRs achieved values parallel to MRs.

### 3.2 Maternal Dependency of Daughter Ramets Under Stolon Severance, Root Restriction, and Drought Stress

#### a) Stolon Severance experiment

##### Relative Water Content (RWC) and survival rate

The relative water content (RWC) of DRs excised from MRs and transplanted into independent pots denoted a clear dependence on developmental stage and duration post-transplantation (Figure 8a). After 8days of separating S1 and S2 DRs from their MRs, RWC dropped significantly (p < 0.05) and reached below 15%. At S3 phase, although DRs experienced a progressive decline in RWC post-transplantation, their water status was better maintained compared to S1 and S2. By day 8, RWC in S3 DRs remained at approximately 53%, which was significantly higher than that of earlier stages, though still substantially lower than the attached MRs. Notably S4 DRs displayed a strikingly different pattern. Their RWC remained relatively stable across the 8-day period, with no significant difference compared to their MRs.

**Figure 8.**
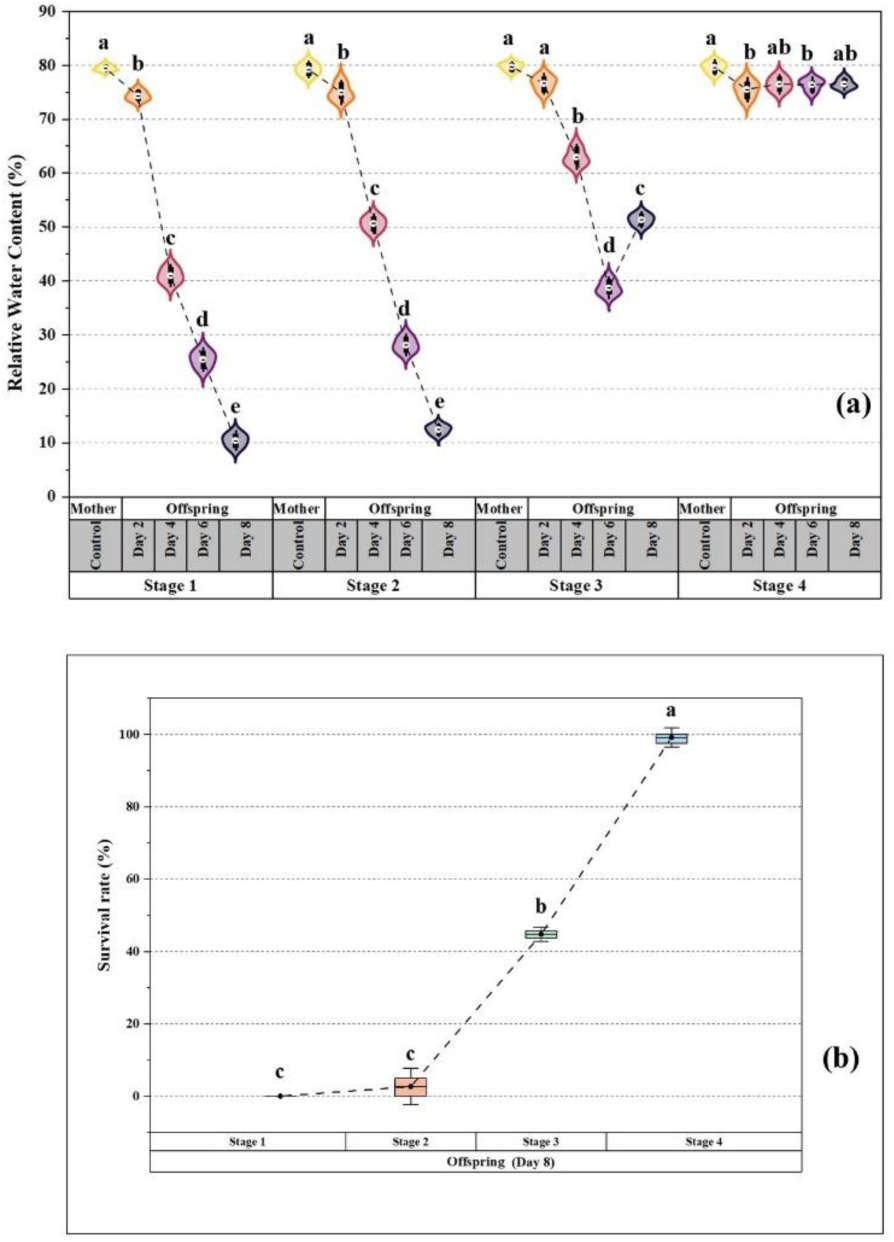
Effect on daughter ramets (DRs) of *Chlorophytum comosum* when separated from mother ramets (MRs) during sequential developmental stages (S1–S4). (a) shows relative water content (RWC), and (b) depicts survival rate of DRs following the cut experiment. Different letters indicate significant differences at *p* ≤ 0.05.

The survival assay conducted on day 8 further emphasized this developmental stage-dependent ability of DRs to establish independently (Figure 8b). DRs at S1 and S2 exhibited negligible survival (0–5%), whereas S3 offspring achieved a moderate survival rate of 45%. Remarkably, S4 DRs showed the highest survival (95%), which was significantly greater than all other stages (Fig. 8b).

#### **b)** Root prevention experiment (S3 stage)

##### Stolon anatomy

After 16 days of preventing root establishment (Stage 3a, 34 days old), stolons remained green and intact, suggesting that functional connectivity between MRs and DRs was maintained despite the absence of daughter roots. In comparison, the normal developmental stage (S5, 34 days old) showed a slight browning of stolons, indicating the early onset of senescence under normal conditions. At S3a phase, where root establishment of DRs was prevented, transverse sections of stolons revealed an intact epidermal layer with no visible degradation. The cortex appeared compact, and vascular tissues (xylem and phloem) remained continuous, showing no signs of collapse. Even after 16 days (similar to S5 age group), the stolon maintained structural integrity confirming that anatomical connectivity was preserved between MRs and DRs despite the absence of root penetration (Fig. 9a).

**Figure 9.**
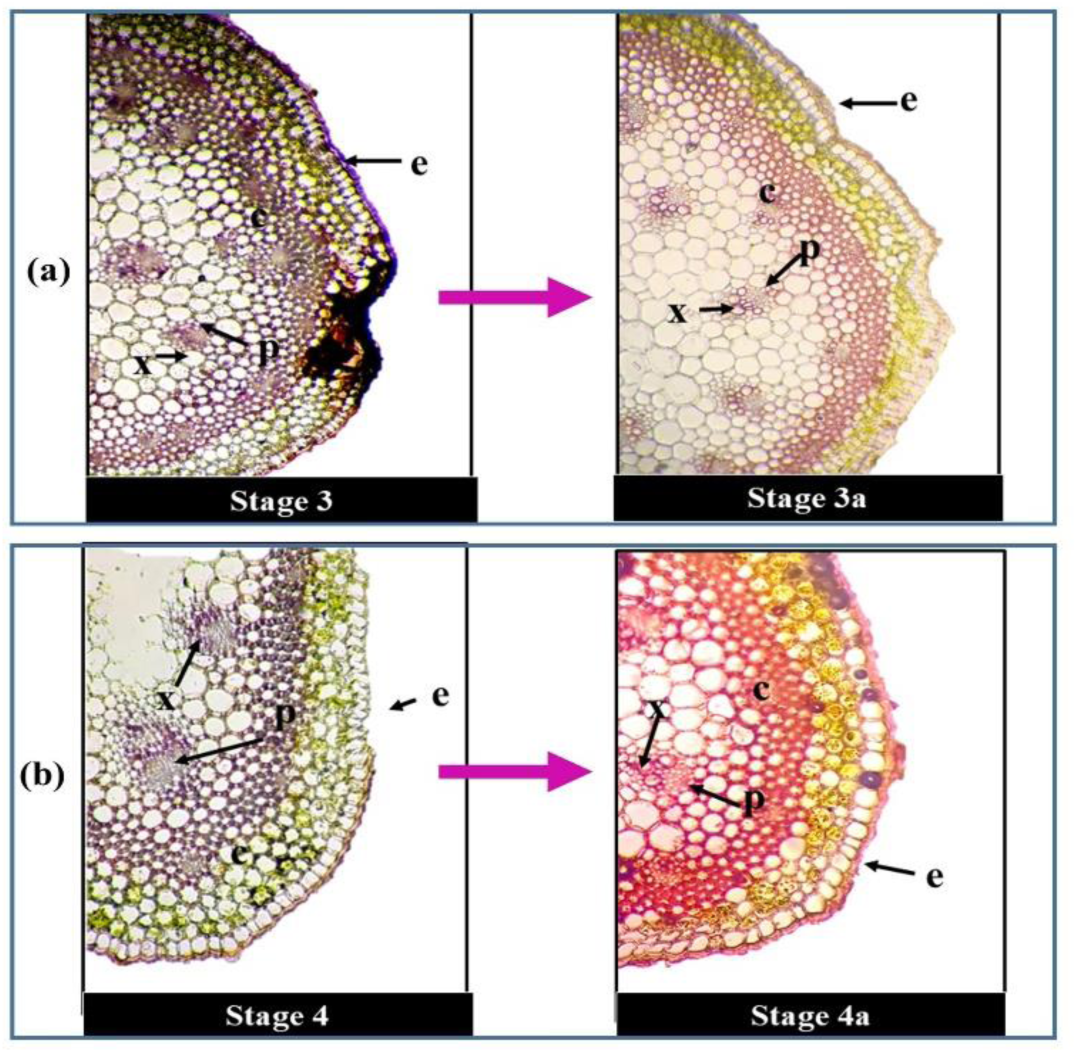
Photomicrographic illustration of transverse sections of stolons in *Chlorophytum comosum* subjected to (a) root prevention experiment and (b) drought-induced experiment. The sections highlight tissue differentiation with distinct epidermis (e), collenchyma (c), chlorenchyma (ch), phloem (p), and xylem (x), representing the organization of different tissue elements under the two experimental conditions.

##### Chlorophyll *a* fluorescence kinetics and JIP test parameters

In the root prevention experiment, chlorophyll *a* fluorescence parameters of MRs and DRs were assessed after 16 days of root establishment inhibition, corresponding to 34-day-old plants. DRs at the S3a stage exhibited markedly reduced photosynthetic efficiency compared with their associated MRs. Specifically, in DRs, F_m_ declined by 18%, φPo by 19%, φEo by 37%, and PIcs by 81%. Moreover, DRs showed a 50% reduction in RC/CS, 33% in TR/CS, and 49% in ET/CS, whereas DI/CS increased by 45% relative to MRs, indicating higher energy dissipation. When MRs of the root prevention set (S3a, 34 days) were compared with naturally developing MRs (S5, 34 days), photosynthetic competence was enhanced under stress. MRs of the root prevention set exhibited a 25% higher PIcs, a 52% increase in RC/CS, and a 20% higher ET/CS. Concomitantly, DI/CS decreased by 18%, while φPo and φEo were elevated by 7% and 17%, respectively (Table 1). These findings suggest that MRs compensated for the impaired root development of DRs by enhancing their photosynthetic performance, thereby maintaining overall physiological homeostasis under stress.

#### **c)** Drought induction experiment (S4 stage)

##### Stolon morphology and anatomy

At S4a phase, where DRs were subjected to drought stress, stolon transverse sections showed cortical shrinkage and slight disorganization in phloem tissues. However, the vascular bundles remained intact, and the epidermis exhibited thickened walls with indications of lignification. Despite external browning, the stolon retained structural continuity, suggesting that functional integrity of vascular tissues was maintained (Fig. 9b). This preserved connection indicates that DRs, unable to extract sufficient water from soil under stress, continued to rely anatomically on stolon-mediated support from the MRs. After 8 days of water deprivation to DRs (S4a, 34 days old), stolons remained green and intact, demonstrating that mother-to-daughter water support preserved stolon integrity under drought conditions. In normal control conditions (S5, 34 days old), stolons showed slight browning and degradation of vascular tissues.

##### Chlorophyll *a* fluorescence kinetics and JIP test

In the drought induction experiment, drought stress was introduced in DRs of S4 development phase when fully established in soil. After 8 days of drought induction, JIP test parameters were assessed in MRs and DRs (S4a, 34 days old). DRs exposed to drought stress demonstrated a pronounced decline in photosynthetic efficiency compared with their corresponding MRs. Specifically, in DRs (S4a), F_m_ decreased by 17%, φPo by 51%, φEo by, and RC/CS up to79%. Similarly, ET/CS declined by 75%, and PIcs was reduced by 88%. Conversely, DI/CS more than doubled in DRs, showing a 143% increase, indicating higher dissipation of absorbed light energy under stress. In contrast, MRs (S4a) enhanced photosynthetic performance compared with both their stressed DRs and naturally developing MRs at the S5 stage (34 days old). Compared with MRs S5, MRs S4a showed increases of 8% in F_m_, 5% in φPo, and 21% in φEo. RC/CS was elevated by 33%, ET/CS by 21%, and PIcs more than doubled (Table 1). Meanwhile, DI/CS decreased by 16%, indicating lower energy loss as heat.

## 4. Discussion

Maternal care is a pervasive and evolutionarily conserved trait across the animal kingdom, commencing at zygote formation and continuing throughout various developmental phases. In mammals, the placenta and umbilical cord serve as critical channels for transporting water and nutrients, which are essential for embryonic development and survival [15,16]. Higher plants exhibit analogous structures, where seeds are sustained by the maternal tissue via the placental region of the ovary until dispersal [17,18]. Despite these parallels, explicit evidence of post-seed maternal care in plants has remained undocumented. To address this gap, we used *Chlorophytum comosum* (spider plant) as a model system, positing that its stolon-mediated clonal propagation functions as a physiological analogue of maternal care, directly supporting the development of its DRs.

### Maternal Care Dynamics Across Sequential Developmental Stages of Daughter Ramets

The anatomical progression of the stolon in *C. comosum* mirrors a maternal care strategy in plants, whereby structural differentiation ensures effective sustenance of developing ramets. In early stages (S1–S2), the thin epidermis, poorly defined procambium, and limited vascular development reflect a sink-dominant state, fully reliant on maternal supply. The gradual thickening of cortex and vascular differentiation by S3–S4 represents the establishment of efficient transport channels, analogous to the functional maturation of umbilical or placental tissues in animals, securing nutrient and water delivery until the daughter achieves autonomy. The onset of senescence at S5–S6, marked by cortical collapse and vascular disintegration, coincides with the attainment of metabolic independence by DRs, indicating a deliberate withdrawal of maternal investment once DRs survival is assured. To the best of our knowledge, this is the first explicit anatomical evidence demonstrating the temporal coordination of vascular differentiation and degeneration in stolons, highlighting their role as transient maternal lifelines that sustain developing ramets until metabolic independence is achieved.

RuBisCO is the most copious protein on Earth, central enzyme of the Calvin–Benson–Bassham cycle and the primary determinant of leaf carboxylation capacity, exhibited a pronounced developmental gradient in DRs maintained under stolon attachment [19–21]. During the early developmental stages (S1–S2), DRs exhibited strongly suppressed RuBisCO activity relative to MRs, reflecting their limited intrinsic capacity for CO₂ fixation and complete dependence on maternally supplied photoassimilates via stolons [22]. This developmental pattern aligns with observations in other species, where the lowest RuBisCO activity has been reported in young, developing mango and olive leaves, while fully expanded mature leaves exhibit substantially higher activity [23,24]. Parallel to this, SPS activity was also minimal, indicating a restricted potential for sucrose biosynthesis and export, which aligns with the severely depressed RuBisCO activity at these stages. As DRs progressed to S3, both RuBisCO and SPS activities showed partial recovery, reflecting the gradual assembly of the photosynthetic apparatus and the onset of autonomous carbon assimilation. By stage S4, enzymatic activities in DRs reached levels comparable to MRs, marking a functional transition to full photosynthetic and carbon autonomy. This developmental pattern is consistent with observations in other species such as sugar beet and *Hevea brasiliensis*, young expanding leaves displayed approximately 2.5-fold lower SPS activity than mature leaves, highlighting their role as sink tissues [25,26].

Acid invertase activity followed a reciprocal trajectory between MRs and DRs, highlighting complementary roles in carbon metabolism during clonal development. DRs exhibited exceptionally high activity at S1, facilitating sucrose hydrolysis to provide hexoses for respiration and biosynthesis when RuBisCO activity was minimal. This strong sink demand declined progressively through S2–S4, converging with maternal levels as DRs gained metabolic independence. Similar trends have been observed in young developing leaves of *Solanum melongena*, *Manihot esculenta*, *Vitis vinifera*, and *Hevea brasiliensis*, where high invertase activity correlates with strong sucrose import and sink strength during growth [26,27]. In contrast, MRs began with low activity but showed a gradual increase, supporting long-distance carbohydrate mobilization. Such reciprocal dynamics indicate a coordinated shift in sink–source balance, as DRs matured, reliance on maternal buffering diminished with the establishment of autonomous photosynthesis and sucrose biosynthesis.

The contrasting NR activity and starch accumulation profiles between MRs and DRs underscore coordinated developmental regulation of nitrogen and carbon metabolism during clonal establishment. MRs maintained consistently high NR activity and stable starch content across stages, serving as reliable sources of reduced nitrogen and carbon to support dependent DRs. In early DR stages (S1–S3), negligible NR activity and very low starch levels reflected the absence of functional roots, limited photosynthetic capacity, and complete reliance on maternal supply, similar to observations in maize seedlings [28]. Progressive increases in NR activity and starch accumulation at S2–S3 indicate the establishment of photosynthetic machinery and onset of autonomous carbon and nitrogen assimilation. By S4, DRs achieved NR activity and starch content approaching maternal levels, marking near-complete metabolic independence. These findings reveal tightly coordinated source–sink dynamics, where MRs buffer early metabolic demand, and DRs progressively assume self-sufficiency through synchronized activation of photosynthetic and assimilatory pathways. Consistent with previous reports, young expanding leaves exhibit low photosynthetic capacity due to delayed chloroplast development, reduced starch accumulation, and limited activity of photosynthetic enzymes [29,30].

The OJIP transients of MRs exhibited a characteristic polyphasic rise, denoting intact and highly efficient PSII performance. In contrast, the markedly flattened transients in S1 DRs, with lower O, J, I, and P steps, indicate photochemical incompetence and disorganization of PSII–LHCII complexes [31]. The reduction in the initial fluorescence (Fo) of S1 DRs suggests a loss of functional antenna connectivity to active PSII reaction centers (RCs), likely reflecting a partial inactivation or structural disorganization of PSII–LHCII complexes [32]. Concomitantly, the reduced Fm values imply restricted electron flow from the primary quinone acceptor (Q_A_) to the plastoquinone pool, which is often symptomatic of limited D1 protein turnover and an imbalance in the redox poise of Q_A_/Q_A_⁻ [13,33,34]. Such photochemical impairments are characteristic of tissues with underdeveloped thylakoid architecture and incomplete chloroplast biogenesis. The gradual restoration of I and P steps through S2 to S4 phases signifies activation of the photosynthetic apparatus, consistent with the reassembly of thylakoid membranes, chloroplast biogenesis, and improved stromal electron transport capacity during clonal establishment.

The developmental progression in RC/CS provides strong quantitative evidence of PSII reorganization during clonal independence. The drastically reduced RC density in S1 DRs likely reflects limited synthesis of core D1/D2 proteins and incomplete oxygen-evolving complex formation. The compensatory overshoot in RC/CS at S4, surpassing maternal values, signifies an adaptive upregulation of PSII density to optimize light capture under emerging autonomous conditions. This mirrors findings in developing leaves of higher plants where chloroplast proliferation peaks during the transition from sink to source status. The initial elevation of ABS/RC and TR_0_/RC in S1 DRs likely reflects a compensatory increase in the excitation energy load per functional RC. Under conditions of limited PSII connectivity and reduced RC density, each active RC must absorb and trap a higher quantum of photons to maintain minimal photochemical activity [36]. However, this elevated excitation pressure induces overreduction of Q_A_ and enhance the probability of energy dissipation as heat. The high DI_0_/RC values in S1 DRs corroborate this photoprotective dissipation mechanism, serving to mitigate photodamage under suboptimal developmental conditions. As thylakoid organization improves during S2–S3 transitions, the ABS/RC and TR/RC values normalize, indicating the recruitment of additional RCs and a reduction in excitation pressure per RC. The stabilization of ET/RC across stages reflects the resilience of downstream electron carriers, suggesting that the core electron transport capacity remains inherently stable once excitation energy is efficiently trapped [37].

At the phenomenological level, the initially depressed ABS/CS, TR/CS, and ET/CS in S1 DRs imply a restricted cross-sectional capacity for light harvesting and energy conversion, possibly due to low chlorophyll density and incomplete expansion of the photosynthetic lamellae [38]. The progressive increase in these parameters across developmental stages parallels the structural expansion of photosynthetic tissue, establishment of intercellular connectivity, and chloroplast proliferation. Interestingly, DI/CS remained low in early DRs despite high DI/RC, suggesting that the total energy dissipated per unit leaf area was constrained by the limited number of active PSII units.

The trends in quantum yield parameters further reinforce the staged recovery of PSII function. The sharp decline of ΦPo and ΦEo in S1 DRs indicates impaired charge separation and electron transport beyond Q_A_, respectively, consistent with the high ABS/RC and DI_0_/RC [39]. The gradual normalization of these parameters through S2–S4 reflects improved energy partitioning between photochemical and non-photochemical pathways, enhanced Q_A_ reoxidation, and a balanced PSII–PSI electron transfer chain. This sequential recovery pattern underscores the tight developmental regulation of photosynthetic efficiency, wherein reactivation of PSII precedes full restoration of downstream carbon metabolism. The performance index (PIcs) serves as a sensitive integrative indicator of the overall photochemical performance, encompassing energy absorption, trapping, and electron transport. The drastic collapse of PIcs in S1 DRs underscores a near-complete functional impairment of PSII during early development, likely due to oxidative stress, low nitrogen assimilation, and insufficient biosynthesis of photosynthetic pigments and proteins. The dramatic increase in PIcs at S4 stages suggests that once the structural and biochemical prerequisites are fulfilled, DRs can even surpass maternal efficiency, reflecting a rejuvenation-like phenomenon typical of newly formed photosynthetic tissues with optimal chloroplast redox homeostasis and efficient excitation energy partitioning.

### Maternal Dependency of Daughter Ramets Under Stolon Severance, Root Restriction, and Drought Stress

#### Stolon Severance experiment

The cut experiment revealed a distinct stage-dependent variation in the relative water content (RWC) of DRs following detachment from the MRs. The RWC of DRs after eight days of detachment from MRs revealed a clear stage-dependent response. During the early developmental stages (S1 and S2), the decline in RWC was severe and irreversible, reflecting the inability of young ramets to maintain water homeostasis once maternal support was removed. In contrast, DRs detached at later developmental stages (S3 and S4) exhibited a markedly improved capacity to establish themselves in soil, maintaining significantly higher RWC values. This physiological trend was mirrored in survival outcomes: while early-stage DRs were unable to survive independently, their drastic decline in RWC to below 15% by day 8 and negligible survival demonstrate that maternal connections are indispensable during this vulnerable period. The intermediate performance of S3 DRs, which sustained moderate RWC (53%) and achieved a survival rate of 45%, suggests a transitional phase of partial autonomy.

At this stage, maternal support still enhances resilience, but DRs begin to develop sufficient physiological competence to tolerate stress in its absence. While, later-stage S4 DRs displayed near-complete independence, with stable RWC comparable to the MRs and a high survival rate. This developmental progression demonstrated substantial survival, highlighting a developmental threshold at which DRs transition from complete maternal dependence to autonomous establishment [36, 38].

#### Root prevention experiment (S3 stage)

The root prevention experiment revealed that stolon-mediated integration ensures functional continuity between MRs and DRs even when root development is inhibited. The maintenance of green, structurally intact stolons with continuous vascular bundles and compact cortex indicates sustained xylem–phloem connectivity, enabling efficient long-distance transport of water, carbon, and signalling metabolites. This structural persistence suggests that hydraulic and assimilate fluxes through the stolon compensate for the absence of daughter roots, thereby maintaining tissue hydration and delaying senescence. The lack of cortical degradation or vascular collapse demonstrates that stolon tissues remain metabolically active, acting as a physiological bridge that safeguards system-level homeostasis under root deprivation. Chlorophyll *a* fluorescence kinetics corroborated this compensatory mechanism. DRs exhibited strong photochemical inhibition, as reflected by reduced F_m_, φPo, φEo, and PIcs, whereas MRs showed enhanced PIcs, RC/CS, and ET/CS alongside decreased DI/CS, signifying an upregulated photosynthetic electron transport chain. This differential response implies that MRs increase PSII performance to offset the impaired carbon fixation of DRs, supporting a unidirectional flow of resources through stolon vasculature [39,40,42,43]. Such modulation of photosynthetic capacity and sustained stolon conductivity exemplify a clonal-level adaptive mechanism akin to maternal care, where physiological plasticity of the MRs ensures the survival and metabolic stability of dependent DRs under adverse or developmentally constrained conditions.

### Drought induction experiment (S4 stage)

The drought induction experiment demonstrated that MRs maintain physiological and structural connectivity to buffer drought-stressed DRs. Despite visible cortical shrinkage and partial disorganization in phloem tissue, the stolons retained intact vascular bundles and a thickened, lignified epidermis, indicating mechanical reinforcement against dehydration. This persistence of vascular continuity suggests that water and assimilate transport from MRs to DRs remained operational, enabling hydraulic compensation when DR roots failed to extract adequate moisture. The absence of complete vascular collapse implies that stolons act as capillary conduits sustaining minimal hydration and metabolite exchange, thereby preserving systemic integrity under drought.

Chlorophyll *a* fluorescence kinetics corroborate this interpretation. The pronounced declines in F_m_, φPo, φEo, RC/CS, ET/CS, and PIcs in drought-stressed DRs reflect severe inhibition of PSII reaction centers, reduced electron transport efficiency, and impaired photochemical energy conversion, likely resulting from thylakoid membrane damage and limited stromal electron acceptors under water deficit. The sharp increase in DI/CS (143%) indicates a photoprotective shift toward non-photochemical quenching, minimizing oxidative damage in dehydrated tissues [36]. In contrast, MRs (S4a) enhanced their PSII functionality, as evidenced by higher F_m_, φPo, φEo, RC/CS, ET/CS, and PIcs, accompanied by reduced DI/CS. This compensatory enhancement suggests an upregulation of photosynthetic electron transport and excitation energy trapping, enabling MRs to increase carbon assimilation and energy flux toward dependent DRs [43–46]. Such differential regulation between MRs and DRs exemplifies a physiological maternal care mechanism, where MRs actively adjust their photosynthetic machinery to sustain clonal integration and metabolic stability under adverse conditions.

## 5. Conclusion

This study provides compelling evidence that stolon-mediated integration in *Chlorophytum comosum* functions as a plant analogue of maternal care, ensuring the survival and metabolic stability of developing daughter ramets (DRs). Anatomical, biochemical, and physiological analyses collectively demonstrate that stolons serve as transient physiological lifelines—facilitating nutrient, water, and signal transport until the DRs attain metabolic autonomy. Early-stage DRs exhibited strong sink behavior, characterized by low RuBisCO and SPS activities, high invertase activity, minimal NR activity, and impaired PSII performance, reflecting complete dependence on maternal support. Progressive enhancement of photosynthetic enzymes, nitrogen assimilation, and PSII efficiency through successive stages (S2–S4) marked the transition to self-sufficiency. Experimental severance, root restriction, and drought assays further revealed that maternal ramets (MRs) dynamically modulate their photosynthetic performance and resource allocation to buffer developing or stressed DRs. The maintenance of stolon vascular continuity and compensatory enhancement of MR photosynthetic activity under adverse conditions highlight a clonal-level adaptive mechanism of parental investment.

Collectively, these findings highlights that even after becoming independent at S4, DRs remain attached to the MRs for a few additional days. If, during this period, the DRs encounters any environmental stress, the MRs exhibits maternal care by re-establishing resource allocation through the stolon-supplying water, nutrients, and other essential metabolites to support the DRs recovery and survival. This experiment on *C. comosum* clearly reveals the maternal care in plants.

## 6. Limitations of the Study

The present study provides the first experimental evidence of maternal care behavior in plants, demonstrated through sustained physiological connections between mother and daughter ramets of *Chlorophytum comosum*. However, as this discovery was made in a clonally propagated species, further research is required to determine whether such maternal mechanisms operate in sexually reproducing and seed-grown plants. Moreover, the experiments were conducted under controlled environmental conditions; therefore, to validate and extend these findings, field-based investigations are strongly recommended to assess maternal interactions under natural ecological dynamics. Additionally, this study examined maternal care under a limited range of stress conditions. Future work should explore a wider spectrum of abiotic and biotic stresses, including nutrient limitation, temperature extremes, heavy metal exposure, and herbivory, to elucidate the broader ecological and adaptive significance of maternal care in plants.

## Consent for Publication

All authors have given their consent for publication.

## Data Availability Statement

The datasets generated and analyzed during the current study are available from the corresponding author upon reasonable request.

## Acknowledgements

The authors sincerely acknowledge Prof. P.L. Swarnkar, Prof. Reto J. Strasser, and Prof. Govindjee for their invaluable guidance and support throughout the study.

## Authors’ Contributions

UB and YS performed the experiments. VS conceived the idea, conceptualized the research, supervised the study, and provided overall guidance. UB and YS drafted the manuscript and prepared the figures, while VS critically reviewed and finalized the manuscript.

