## Supplementary material for "Biochemical and Physiological Evidence of Maternal Care in Plants: A Case Study of *Chlorophytum comosum*": 2 figures and 1 table

### Supplimentry materials

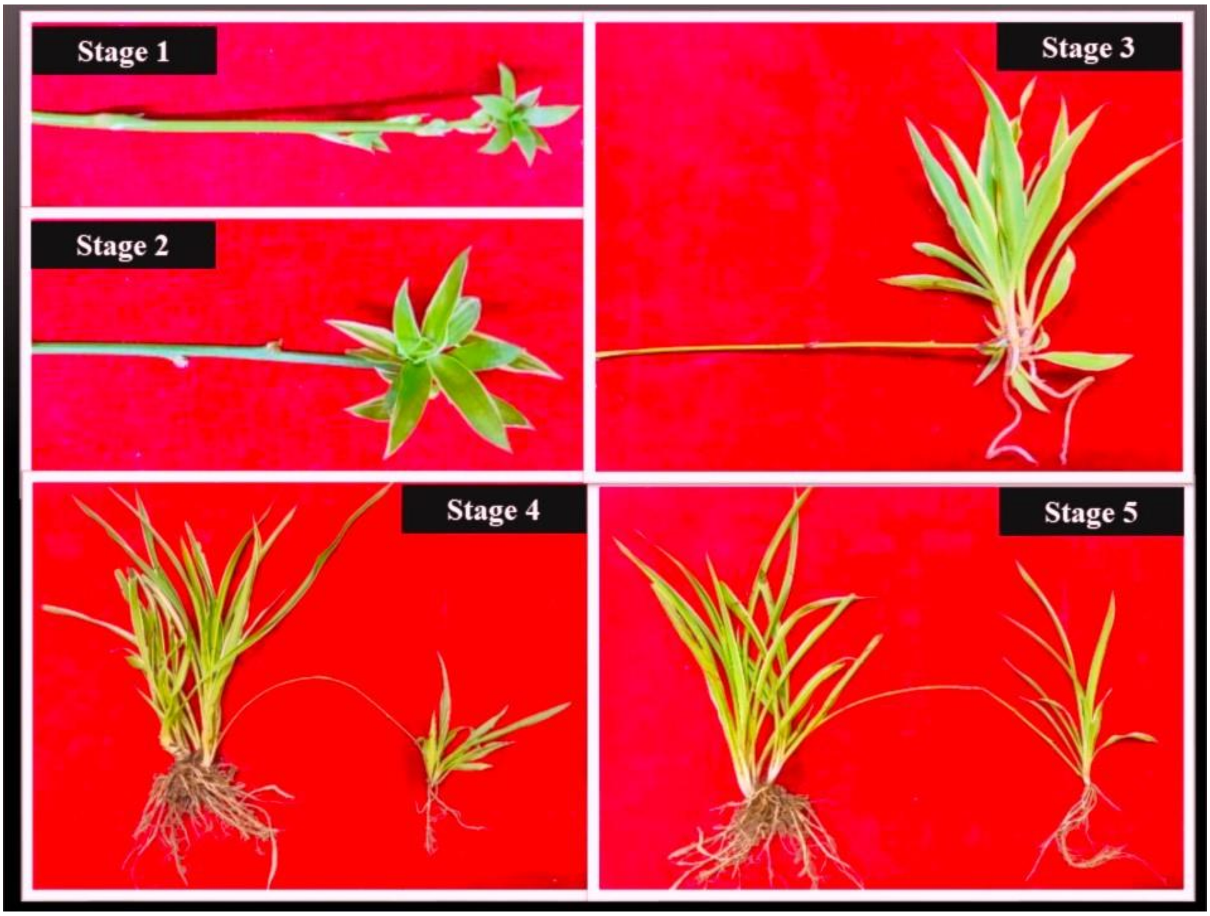


**SF Figure 1:** *Sequential developmental stages of* daughter *ramets (DRs) of Chlorophytum comosum showing their physiological connection with the mother ramet (MR) through stolonal linkage*. The images represent original photographs depicting the progressive ontogeny of *DRs* at distinct developmental stages under controlled growth conditions.

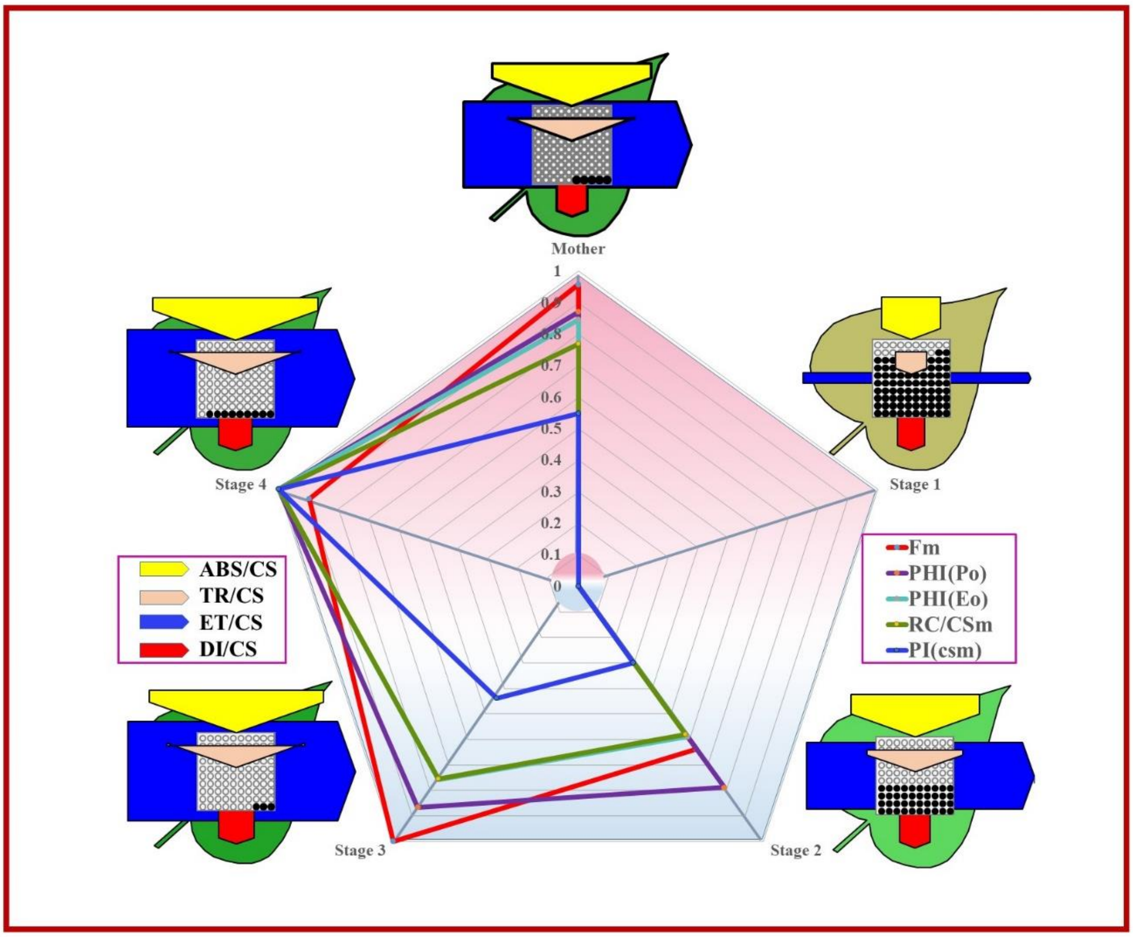


**SF Figure 2:** *Comparative phenomenological flux model and radar plot depicting photosynthetic performance of Chlorophytum comosum mother ramets (MRs) and daughter ramets (DRs) across four developmental phases (S1–S4)*. The phenomenological flux model illustrates the energy fluxes per excited cross-section (ABS/CS, TR/CS, ET/CS, and DI/CS), representing light absorption, energy trapping, electron transport, and energy dissipation, respectively. The accompanying radar plot quantitatively compares these flux parameters between MRs and DRs at each developmental stage, highlighting phase-dependent modulation of energy partitioning during development.

| Parameter | Definition / Description | Formula |
|---|---|---|
| **Fo** | Minimal fluorescence when all PSII reaction centers (RCs) are open | $F_0 \cong F_{50\mu s}$ |
| **Fm** | Maximal fluorescence when all PSII RCs are closed | $F_m \cong F_P$ |
| **ABS/RC** | Absorption flux per RC | $\frac{ABC}{RC} = M_O.\left(\frac{1}{V_j}\right).\left(\frac{1}{\varphi P_O}\right)$ |
| **TRo/RC** | Trapped energy flux per RC leading to $Q_A$ reduction | $TR/RC = M_o \times (1 -)$ |
| **ETo/RC** | Electron transport flux per RC beyond $Q_A$ | $\frac{ET}{RC} = M_O . \left[1/\frac{F_{2ms} - F_O}{F_M - F_O}\right] . \Psi_O$ |
| **DIo/RC** | Dissipated energy flux per RC | $\frac{DI}{RC} = \left(\frac{ABS}{RC}\right) - [M_O \times \left(\frac{1}{V_j}\right)]$ |
| **ABS/CSm** | Absorption flux per excited cross-section | ABS/CS = Fluorescence intensity at 50µs |
| **TRo/CSm** | Trapped energy flux leading to $Q_A$ reduction per cross-section | $TR/CS = \varphi P_O \cdot (ABS/CS)$ |
| **ETo/CSm** | Electron transport flux beyond $Q_A$ per cross-section | $\frac{ET}{CS} = \varphi E_O. (ABS/CSm)$ |
| **DIo/CSm** | Energy dissipation flux per cross-section | $\frac{DI}{CS} = \left(\frac{ABS}{CS}\right) - [\Phi P_O \times \left(\frac{ABS}{CS}\right)]$ |
| **RC/CSm** | Density of active PSII reaction centers per excited cross-section | $\varphi P_O = \frac{TR}{ABS} = [1 - \left(\frac{F_O}{F_M}\right)]$ |
| **φ(Po)** | Maximum quantum yield of primary photochemistry | $\varphi E_O = \frac{ET}{ABS} = \left[1 - \left(\frac{F_O}{F_M}\right)\right] . \psi_O$ |
| **φ(Eo)** | Quantum yield for electron transport beyond $Q_A$ | ABS/CS = Fluorescence intensity at 50µs |
| **PIcs** | Performance Index on Cross-Section Basis | $PI_{CS} = \frac{ABS}{CS} \times \frac{1 - (F_O/F_M)}{M_O/V_j} \times \frac{F_M - F_O}{F_O} \times \frac{1 - V_j}{V_j}$ |

**SF Table 1:** *Comprehensive summary of JIP-test parameters derived from fast chlorophyll a fluorescence transients (OJIP curve) in Chlorophytum comosum. The table presents original and derived parameters with their mathematical expressions and physiological interpretations.*
